# Distinct cell adhesion signature defines glioblastoma myeloid-derived suppressor cell subsets

**DOI:** 10.1101/2021.09.27.461995

**Authors:** Defne Bayik, Cynthia F. Bartels, Katreya Lovrenert, Dionysios C. Watson, Duo Zhang, Kristen Kay, Adam Lauko, Sadie Johnson, Alice Lo, Mary McGraw, Matthew Grabowski, Alireza M. Mohammadi, Filippo Veglia, Yi Fan, Michael A. Vogelbaum, Peter Scacheri, Justin D. Lathia

**Affiliations:** Lerner Research Institute, Cleveland Clinic, OH; Case Comprehensive Cancer Center, Cleveland, OH; Department of Genetics and Genome Sciences, Case Western Reserve University, Cleveland, OH; University Hospitals Cleveland Medical Center, Cleveland, OH; Department of Radiation Oncology, University of Pennsylvania, Philadelphia, PA; Department of Pathology, Case Western Reserve University, Cleveland, OH; Case Western Reserve University, Medical Science Training Program, Cleveland, OH; Rose Ella Burkhardt Brain Tumor Center, Cleveland Clinic, OH; Department of Immunology, Moffitt Cancer Center, Tampa, FL; Department of Neuro-oncology, Moffitt Cancer Center, Tampa, FL

## Abstract

Increased myeloid-derived suppressor cell (MDSC) frequency is associated with worse outcomes and poor therapeutic response in glioblastoma (GBM). Monocytic (m) MDSCs represent the predominant subset in the GBM microenvironment. However, the molecular basis of mMDSC enrichment in the tumor microenvironment compared to granulocytic (g) MDSCs has yet to be determined. Here, we report that mMDSCs and gMDSCs display differences in their tumor-accelerating ability, with mMDSCs driving tumor growth in GBM models. Epigenetic assessments indicate enhanced gene accessibility for cell adhesion programs in mMDSCs and higher cellsurface integrin expression in mouse and human mMDSCs. Integrin β1 blockage abrogated the tumor-promoting phenotype of mMDSCs and altered the immune profile in the tumor microenvironment. These findings suggest that integrin β1 expression underlies the enrichment of mMDSCs in tumors and represents a putative immunotherapy target to attenuate myeloid cell-driven immune suppression in GBM.

**Summary:** Myeloid-derived suppressor cells (MDSCs) drive glioblastoma growth; however, the function of specific MDSCs subsets is unclear. Bayik *et al.* demonstrate that adhesion programs are enhanced in monocytic MDSCs and responsible for their GBM-promoting function.

## Introduction

Glioblastoma (GBM), the most common primary malignant brain tumor, is characterized by a dramatic infiltration of immunosuppressive myeloid cells, which can comprise 30-50% of the tumor mass (de Groot et al., 2020; De Leo et al., 2020; Gutmann and Kettenmann, 2019; Pinton et al., 2019). Accumulation of these myeloid cells represents a critical barrier to treatment of GBM, and their targeting improves response to radiotherapy and immunotherapy in preclinical models (Antonios et al., 2017; Kamran et al., 2017; Zhang et al., 2019). Together with tumor-associated macrophages and microglia, myeloid-derived suppressor cells (MDSCs) constitute one of the major immunosuppressive myeloid cell populations in GBM (De Leo et al., 2020). Myeloid-derived suppressor cells (MDSCs) are a heterogeneous population of immature cells that originate in the bone marrow (Veglia et al., 2021). Biomarker studies further established that MDSCs expand in the peripheral circulation of patients with GBM compared to those with low-grade brain malignancies, accumulate in tumors and associate with worse disease outcome (Alban et al., 2018; Gielen et al., 2016; Raychaudhuri et al., 2015; Raychaudhuri et al., 2011). These observations have served as the basis for the development and assessment of anti-MDSC therapies in GBM and other advanced cancers (De Cicco et al., 2020; Law et al., 2020; Peereboom et al., 2019).

MDSCs are classified into two phenotypically and functionally distinct subsets, monocytic (mMDSC) and granulocytic/polymorphonuclear (gMDSC) (Veglia et al., 2021). While both subsets can interfere with the activity of cytotoxic T cells, recent studies demonstrate that they undertake additional roles both systemically and within the tumor microenvironment. In breast cancer models, it was demonstrated that mMDSC localization to primary tumors drives a stem cell phenotype, while gMDSCs facilitate metastatic spread to the lungs (Ouzounova et al., 2017). Assessment of MDSC subsets also revealed a difference in their localization in GBM. mMDSCs represented the dominant subtype in both mouse and human tumors and were especially enriched in male tumors (Bayik et al., 2020; Zhang et al., 2019). This was in part driven by differential response to chemoattractants by MDSC subtypes. Both the CCL2-CCR2 and the macrophage migration inhibitory factor (MIF)-CD74 axes have been implicated in the recruitment of mMDSCs to the GBM microenvironment (Alban et al., 2020; Chang et al., 2016; Flores-Toro et al., 2020; Otvos et al., 2016). However, the molecular basis of distinct MDSC subset trafficking and function in the context of GBM remains unclear. We hypothesized that differences in epigenetic architecture between MDSC subsets impacted their trafficking and interaction with the tumor microenvironment and found that cellular adhesion programming, and particularly integrin β1 function, contributed to the distinct pro-tumorigenic function of mMDSCs.

## Results and Discussion

### mMDSCs and gMDSCs have distinct functions and chromatin accessibility signatures

While we previously demonstrated that targeting of mMDSCs but not gMDSCs prolonged survival of male mice with GBM, the inhibitors were nonspecific (Bayik et al., 2020). To more directly interrogate the tumorigenic effects of MDSC subsets on vivo tumor growth, we implanted male mice with syngeneic mouse tumors before adoptively transferring bone marrow-derived mMDSCs or gMDSCs (**Fig. 1A**). The transfer of male mMDSCs reduced the median survival duration by 15-25% in multiple models (both GL261- and SB28-bearing mice), while male gMDSCs had no impact on the median survival span (**Fig. 1B-C**). We previously reported sex differences in MDSC subset activity in GBM, with increased tumor-infiltration of male mMDSCs in animal models and GBM patients (Bayik et al., 2020). Thus, to further explore whether the tumor-promoting effect of mMDSCs was informed by the sex of the cells, we repeated the adoptive transfer experiments using MDSCs isolated from female hosts and observed a similar trend, with mMDSCs accelerating tumorigenesis and gMDSCs having no effect on the course of GBM progression (**Fig. S1A-B**). Collectively, these results suggest that mMDSCs inherently drive progression of GBM in preclinical models to a greater extent than gMDSCs, which show equivalent malignancy to vehicle controls.

**Figure 1:**
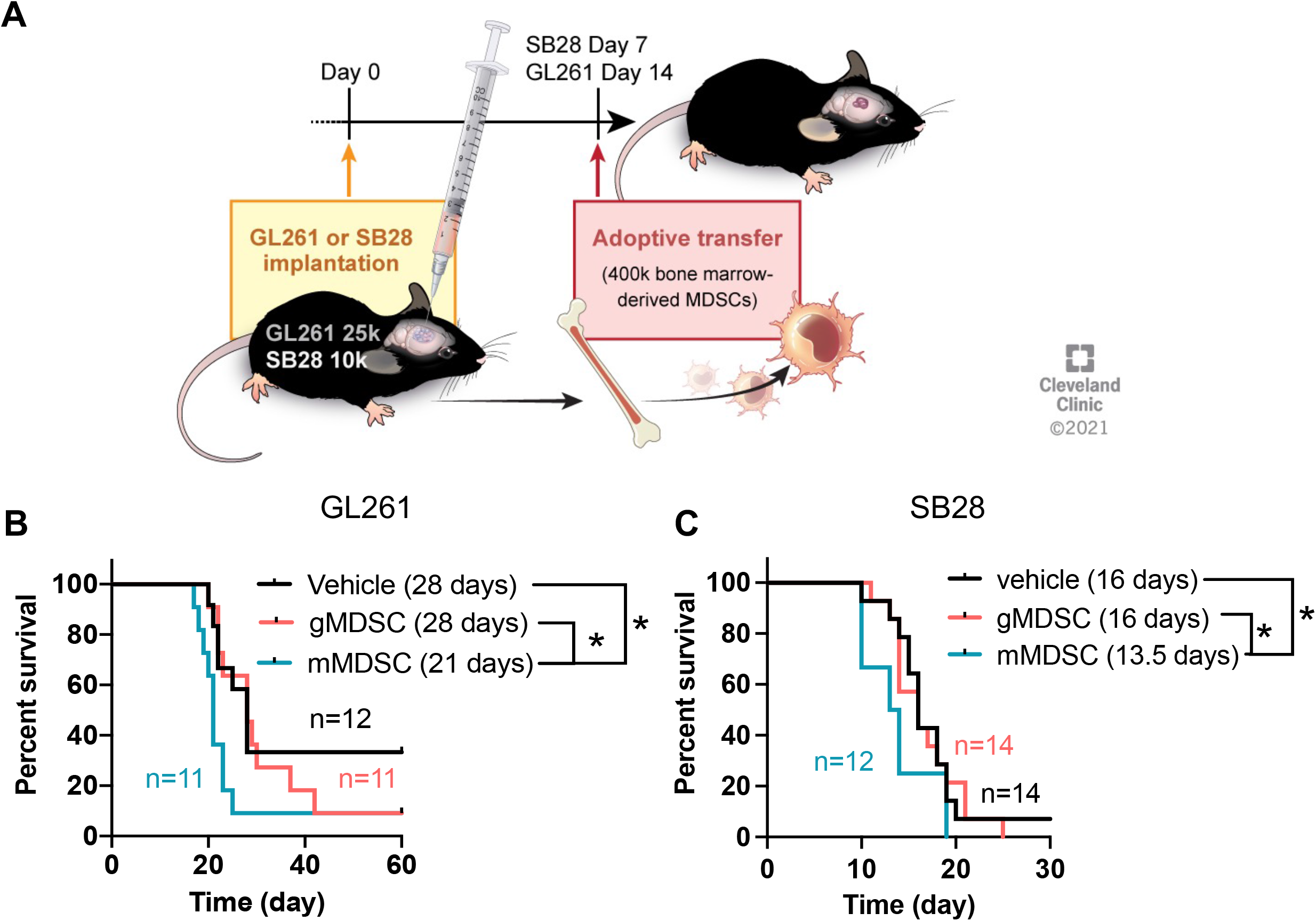
mMDSCs but not gMDSCs promote tumor growth. **(A)** Male C57BL/6 mice were implanted with 25,000 GL261 or 10,000 SB28 cells. Seven (SB28) or 14 (GL261) days post-tumor implantation, mice were adoptively transferred with 400,000 mMDSCs or gMDSCs isolated from the bone marrow of male mice with matching tumors by retro-orbital injection. Kaplan-Meier curves depicting survival of **(B)** GL261- or **(C)** SB28-bearing mice post-adoptive transfer. n=11-14 mice/group from 3 independent experiments. * p<0.05 as assessed by Wilcoxon-rank test.

Several studies have identified unique gene expression signatures associated with MDSC subsets in mouse cancer models and patients with malignancies (Alshetaiwi et al., 2020; Mastio et al., 2019; Sasidharan Nair et al., 2020; Song et al., 2019). However, the epigenetic basis underlying distinct MDSC characteristics and expression profiles remain unclear. Therefore, we performed Assay for Transposase-Accessible Chromatin with high-throughput sequencing (ATAC-seq) on mMDSCs and gMDSCs isolated from the bone marrow of GL261-bearing or control male and female mice (**Fig. 2A**). Unsupervised clustering analyses revealed no significant differences between the chromatin accessibility of MDSCs obtained from sham-versus GL261-injected mice, suggesting that these cells do not undergo major developmental reprogramming in bone marrow in the presence of tumors (**Fig. S1C**). In addition, 27% of the variance in genomewide accessibility was linked to host sex, while subtype identity accounted for 70%, further underscoring the role of cellular identity as the main determinant of the epigenetic architecture of MDSC subsets (**Fig. S1D**). Therefore, we focused on the differentially accessible regions between mMDSCs and gMDSCs by controlling for sex and tumor state. This approach identified a total of >40,000 variably accessible chromatin loci (**Fig. 2B**), of which 12,613 (61%) were gained and 7,929 (39%) were lost peaks in mMDSCs when stratified for peaks with mean count >50. We next performed gene ontology analysis using the peaks gained in mMDSCs to evaluate whether genes localized at these more accessible regions are enriched in specific pathways. This approach revealed “regulation of leukocyte cell-cell adhesion” as the top pathway associated with mMDSCs over gMDSCs (**Fig. 2C**). Complementary to our observations, two recent studies demonstrated that histone deacetylase and DNA methyltransferase inhibitors can interfere with MDSC chemotaxis (Lu et al., 2020; Sasidharan Nair et al., 2020). Together, these observations suggest a mechanism through which cell adhesion and migration programs are epigenetically regulated.

**Figure 2:**
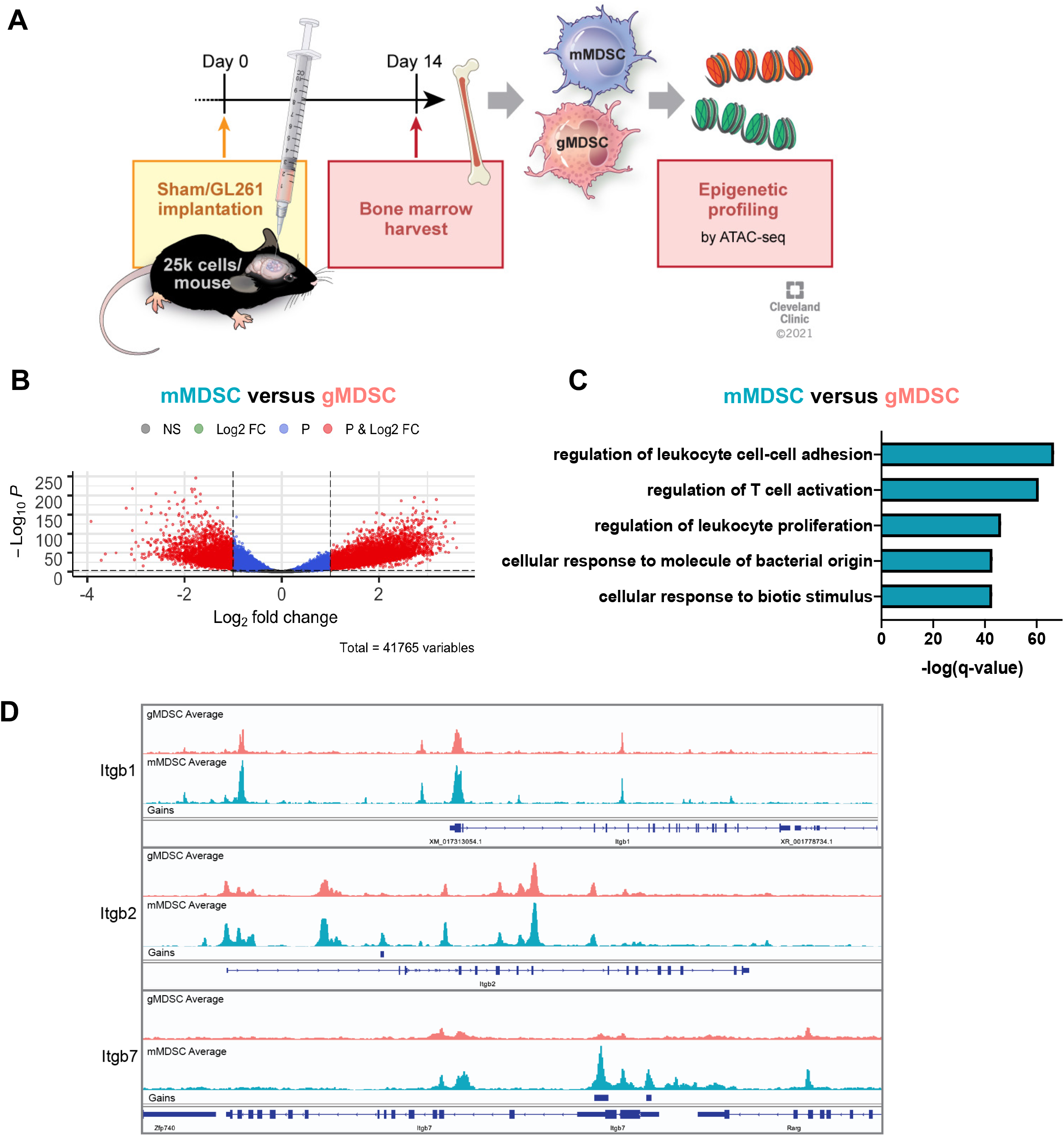
mMDSCs and gMDSCs have distinct epigenetic programming. **(A)** C57BL/6 mice were implanted with 25,000 (male) and 30,000 (female) GL261 cells or were sham injected. Fourteen days later, mMDSCs or gMDSCs were isolated from the bone marrow for ATAC-seq. **(B)** Differentially accessible genes between mMDSCs and gMDSCs with peak count >50 and controlled based on tumor state and sex. **(C)** Top 5 pathways upregulated in mMDSCs with log2 fold change ≥ 1 and adjusted p ≤ 0.001 based on the gained peaks in mMDSCs. **(D)** Cross-section of integrin β1 (Itgb1), integrin β2 (Itgb2) and integrin β7 (Itgb7) loci demonstrating gained peaks in mMDSCs and gMDSCs.

### Integrin β1 is highly expressed in mouse and human mMDSCs

Enhanced adhesion is a common feature of malignant tumors, ranging from being essential for tumor cells to drive the cancer stem cell program, as we have previously demonstrated (Day et al., 2019; Lathia et al., 2010), to being elevated in immune cell populations to drive chemotaxis and tumor infiltration (Harjunpaa et al., 2019). As integrins are a major class of receptors that recognize multiple extracellular matrix (ECM) proteins and are essential for adhesion and downstream signaling (Desgrosellier and Cheresh, 2010), we focused on the potential link between integrins and the elevated adhesion signature in mMDSCs. Integrins are heterodimers formed by interaction of α- and β-chains and show variable expression across cell populations, with integrins β1, β2 and β7 playing a central role in leukocyte migration (Desgrosellier and Cheresh, 2010; Harjunpaa et al., 2019). Of these, α4β1 is particularly important for tumor trafficking of myeloid cells, and inhibition of the PI3Kγ-α4 signaling axis was shown to reduce the frequency of bulk MDSCs in lung and pancreatic cancer models (Foubert et al., 2017; Jin et al., 2006; Schmid et al., 2013). However, there is limited knowledge on the differential integrin expression profile of MDSC subsets and how it is linked to their behavior. ATAC-Seq analysis revealed that Itgb1, Itgb2 and Itgb7 contained open-chromatin regions, highlighting the potential for gene transcription in mMDSCs and gMDSCs (**Fig. 2D**). Analysis of surface integrin β1, β2 and β7 subunits in mice with GL261 or SB28 tumors and sham-injected controls demonstrated that myeloid cells in circulation and in tumors had higher levels of integrins β1 and β7 compared to lymphocytes (**Fig. S2A**, *data not shown).* While integrin β2 levels were also higher in blood myeloid populations, tumor-infiltrating adaptive immune cells upregulated this receptor (**Fig. S2A**). A pairwise comparison of the receptor expression on mMDSCs versus gMDSCs showed that mMDSCs had significantly more surface integrins in the bone marrow of tumor-bearing and healthy mice (**Fig. 3A**). While this pattern was retained for integrins β1 and β7 in blood and tumors, gMDSCs in these compartments had similar or higher levels of integrin β2 compared to mMDSCs (**Fig. 3B-C**). Importantly, there were no significant differences in integrin levels between male and female mMDSCs, supporting the observation that epigenetic regulation is primarily influenced by cell identity (**Fig. S1D-E**). To further validate these observations in GBM patients, we analyzed single-cell expression profiles of integrin β1 (ITGB1), β2 (ITGB2) and β7 (ITGB7) in tumorinfiltrating mMDSCs and gMDSCs using publicly available datasets (Caruso et al., 2020). mMDSCs and gMDSCs were distinguished from other myeloid cells using a combination of CD84, CD33, ITGAM (CD11b), CD14, OLR1 (LOX-1), CEACAM8 (CD66) and HLA-DR (**Fig. 3D**, **S2C**). This comparison demonstrated that mMDSCs expressed higher levels of ITGB1 and ITGB7 but not ITGB2 compared to gMDSCs (**Fig. 3E**). We sought to confirm the correlation between RNA and protein levels by measuring surface integrin β1 and β7 levels in patient specimens. mMDSCs circulating in the blood or localizing to tumors had significantly more integrin β1 compared to gMDSCs, whereas integrin β7 intensity was similar between the two subsets (**Fig. 3F-G**). Myeloid-dominant and consistent expression of integrin β1 in the bone marrow, blood and tumor suggests that integrin β1 is a putative target to regulate immunosuppression in GBM.

**Figure 3:**
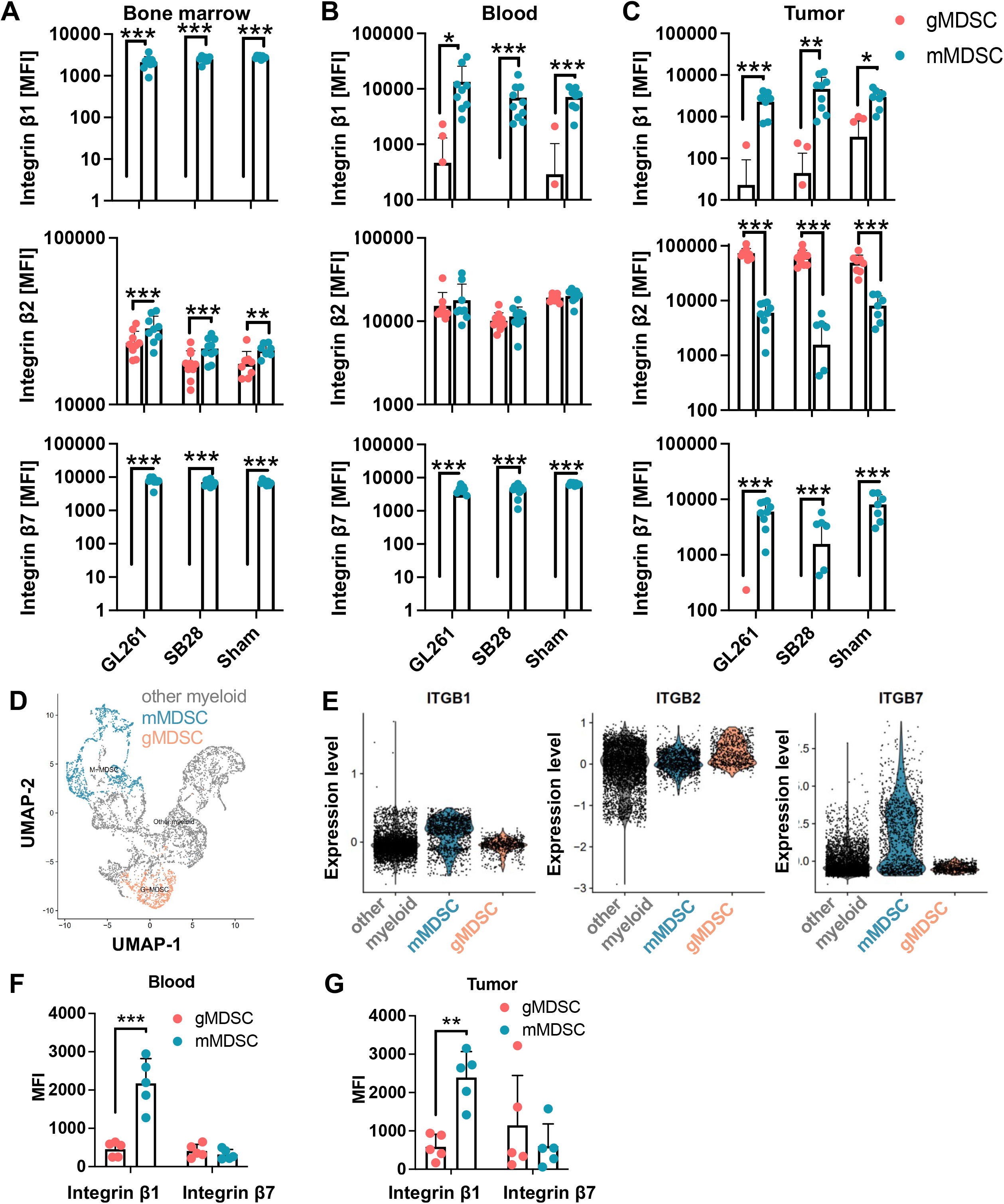
mMDSCs consistently express higher levels of integrin β1. C57BL/6 mice were implanted with 25,000 GL261 cells and 15,000 SB28 cells or sham injected. Mice were euthanized 12 (SB28) or 19 (GL261) days post-tumor implantation. Differential expression of surface integrin β subunits by mouse mMDSCs and gMDSCs localized at **(A)** bone marrow, **(B)** blood and **(C)** tumors/brains. **(D)** UMAP depicting distribution of myeloid cell subpopulations in patient tumors defined based on markers given in Fig. S2B. n=50 combined from Darmanis, Yuan, Neftel and Yu et al. **(E)** Expression levels of integrin β1, β2 and β7 in tumor myeloid cells at a single-cell resolution in myeloid cell populations. Surface integrin β1 and β7 expression on human mMDSCs and gMDSCs from **(F)** blood or **(G)** tumors. n=5 (2 male, 3 female). ** p<0.01 and *** p<0.001 by two-way ANOVA.

### Blockade of integrin β1 abrogates mMDSC function

To evaluate the impact of mMDSC-specific targeting of integrin β1, we pre-treated sorted donor cells with an anti-integrin β1 neutralizing (anti-β1) or control isotype antibody *in vitro* prior to adoptive transfer. Mice that received mMDSCs incubated with anti-β1 antibody had significantly longer tumor latency compared with mice receiving mMDSCs pre-treated with isotype control antibody (p<0.05, **Fig. 4A**). These data indicate that integrin β1 blockade abrogates the tumor-promoting role of these cells. Of note, these differences are likely not due to the induction of MDSC cell death, as we observed no significant difference in cell viability during the 1 hour antibody incubation (*data not shown*). As expected, transfer of donor gMDSCs stimulated with an anti-β1 antibody had no effect on the median survival duration of mice with tumors (**Fig. 4B**). To further assess the specificity of integrin β1 signaling, we repeated the same experiment while instead blocking integrin β7 on mMDSCs. There was no significant difference between the survival span of mice that acquired isotype-versus anti-β7-treated mMDSCs (**Fig. 4C**). Based on this observation that integrin β1 inhibition selectively alters mMDSC function, we focused on the mMDSC-related changes in the tumor microenvironment. Mice were adoptively transferred with mMDSCs and gMDSCs with intact or blocked surface integrin β1, and the frequency of circulating and tumor-infiltrating immune populations was analyzed 3 days later. Overall, leukocyte infiltration, as well as the relative frequency of mMDSC, gMDSCs, monocytes, myeloid DCs, B cells CD4^+^ T cells, CD8^+^ T cells and NK cells, was similar between the isotype and anti-β1 groups (**Fig. S3A-B, Fig. 4D-G**). In contrast, there was a significant increase in the abundance of macrophages with a concomitant reduction in conventional dendritic cell (cDC) frequency in mice with isotype-treated mMDSCs (**Fig. 4H-I**). Importantly, the observed immune changes were limited to the tumor microenvironment and specifically induced by mMDSCs, as gMDSC transfer did not alter the frequency of various innate or adaptive cells in tumors and there were significant variations in peripheral immune populations (**Fig. S3C-D**). Thus, the enhanced macrophage phenotype could be a result of mMDSC polarization into tumor-associated macrophages (Corzo et al., 2010; Kwak et al., 2020) or an indirect consequence of the interaction between mMDSCs and macrophages leading to an accumulation of the latter population (Beury et al., 2014). Blockade of integrin β1 in vitro did not interfere with mMDSC-to-macrophage differentiation or the phenotype of macrophages (**Fig. S3E-F**). These results further underscore that the tumor microenvironment impacts mMDSC fate and support our previous observations that mMDSCs were the predominant subset in the GBM microenvironment (Bayik et al., 2020). While the exact mechanism by which integrin β1 regulates mMDSC-macrophage communication remains to be investigated, our results establish that the cell adhesion machinery is inherently different between MDSC subsets, with mMDSCs having enhanced integrin β1 signaling with functional consequences for anti-tumor immunity (**Fig. 5**). Moreover, these findings support the future development of integrin β1 targeting strategies, particularly as integrin β1 expression correlates with poor GBM outcome and is upregulated in GBM models resistant to anti-angiogenic therapy (Carbonell et al., 2013; Malric et al., 2017). Given that integrin β1 also plays a role in GBM cell proliferation and self-renewal (Carbonell et al., 2013; Malric et al., 2017), such strategies might have the dual benefit of targeting tumor cells while modulating the immune response.

**Figure 4:**
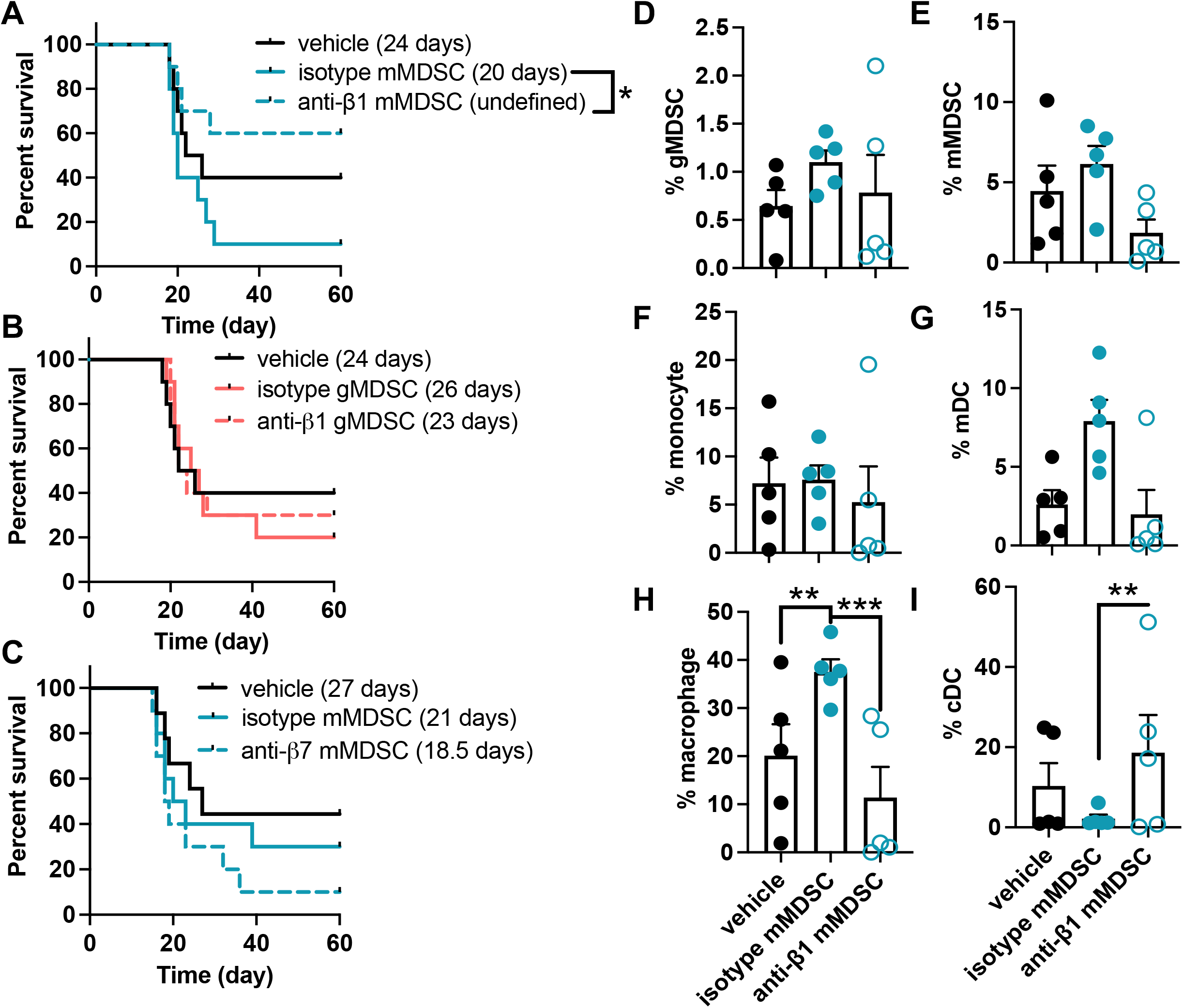
Integrin β1 comprises a therapeutic target to regulate mMDSC function. Bone marrow-derived male mMDSCs and gMDSCs were treated with 100 μg/ml anti-integrin β1, anti-integrin β7 or isotype control antibody prior to adoptive transfer. Kaplan-Meier plot depicting survival of GL261-bearing male mice transferred with **(A)** mMDSCs treated with anti-integrin β1 neutralizing antibody, **(B)** gMDSCs treated with anti-integrin β1 neutralizing antibody or **(C)** mMDSCs treated with anti-integrin β7 neutralizing antibody. n=9-10 from two independent experiments. * p<0.05 as assessed by Wilcoxon-rank test. C57BL/6 male mice were implanted with 25,000 GL261 cells and adoptively transferred with 400,000 mMDSCs treated with isotype control antibody or anti-integrin β1 blocking antibody for 1 hour. Myeloid cell populations were analyzed from tumors 3 days later. The frequency of tumor-infiltrating **(D)** gMDSCs, **(E)** mMDSCs, **(F)** monocytes, **(G)** macrophages, **(H)** myeloid DCs and **(I)** conventional DCs in mice adoptively transferred with isotype- or anti-integrin β1-treated mMDSCs. n=5/group, ** p<0.01 and *** p<0.001 as determined by two-way ANOVA with Tukey’s correction.

**Figure 5:**
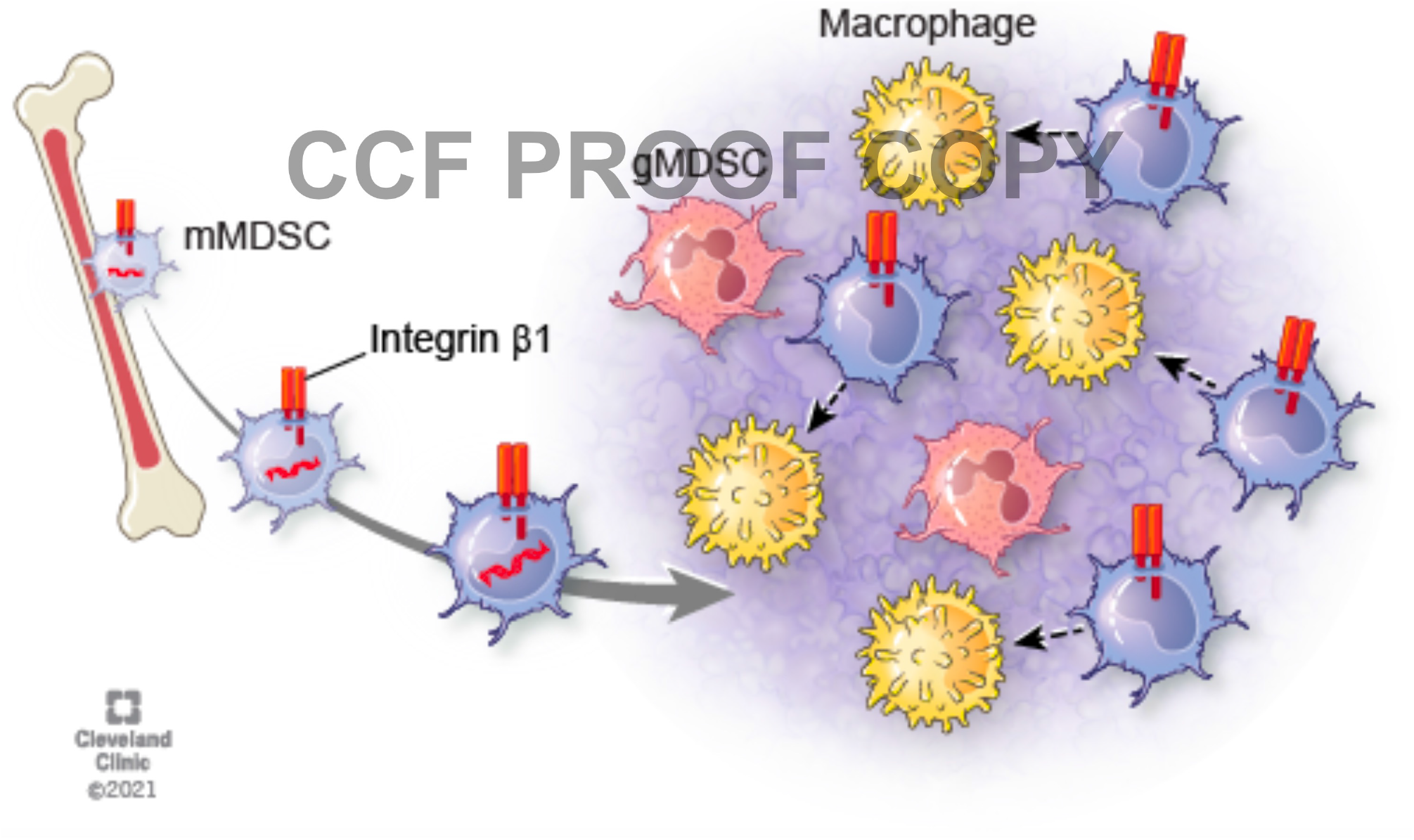
Blockade of integrin β1 expression on mMDSCs reprograms the tumor microenvironment. mMDSCs, which are the abundant MDSC subset in GBM, express high levels of integrin β1. Inhibition of this signaling axis on mMDSCs presented as reduced macrophage accumulation in tumors.

## Acknowledgements

The authors thank Dr. Erin Mulkearns-Hubert for editorial assistance, Ms. Amanda Mendelsohn for illustrative work and Dr. Thad Stappenbeck’s laboratory for equipment access and Lerner Research Institute Flow Cytometer Core for sorting assistance. This work is supported by National Institutes of Health grants R01NS109742 (J.D. Lathia & M.A. Vogelbaum), P01CA245705 (J.D. Lathia), K99CA248611 (D. Bayik), T32AI007024 and TL1TR002549 (D.C. Watson), F31CA264849 (K. Kay), and F30CA250254 (A. Lauko), R01NS094533 (Y. Fan), R01NS106108 (Y. Fan), R01CA241501 (Y. Fan), R01HL155198 (Y. Fan), and American Heart Association Predoctoral Fellowship 830890 (D. Zhang).

## Author contributions

D. Bayik designed the study, performed the experiments, analyzed results, interpreted data and wrote the manuscript; C. Bartels performed the experiments and analyzed results; K. Lovrenert analyzed results and interpreted data; D.C. Watson performed the experiments, analyzed results and interpreted data; D. Zhang performed the experiments and analyzed results, K. Kay performed the experiments; A. Lauko performed the experiments, and interpreted data; S. Johnson performed the experiments; A. Lo performed the experiments; M. McGraw generated critical samples; M. Grabowski generated critical samples; A.M. Mohammadi generated critical samples; F. Veglia interpreted data; Y. Fan supervised the experiments; M.A. Vogelbaum supervised the experiments; Peter Scacheri interpreted data and supervised the experiments; J.D. Lathia designed the study, interpreted data, supervised the experiments and wrote the manuscript. All authors read and approved the final manuscript.

## Methods

### Cell lines

GL261 cells were obtained from the Developmental Therapeutics Program, National Cancer Institute, and the SB28 line was gifted by Dr. Hideho Okada (University of California, San Francisco). Cells were maintained in RPMI 1640 (Media Preparation Core, Cleveland Clinic) supplemented with 10% FBS (Thermo Fisher Scientific) and 1% penicillin/streptomycin (1% Pen/Strep, Media Preparation Core). All cell lines were treated with 1:100 MycoRemoval Agent (MP Biomedicals) upon thawing and routinely tested for Mycoplasma spp. (Lonza).

### Antibodies

For sorting and immune profiling, the following antibodies were purchased from Biolegend, unless otherwise specified: Gr-1 (clone RB6-8C5, 11-5931-85, eBioscience), CD11b (clone M1/70, 563553, BD Bioscience), CD11c (clone HL3, 612796, BD Bioscience), Ly6G (clone 1A8, 560603, BD Bioscience), CD3 (clone 145-2C11, 56379, BD Biosciences), γδ TCR (clone GL3, 562892, BD Bioscience), I-A/I-E (clone M5/114.15.2, 107606), CD68 (clone FA-11, 137024), Ly6C (HK1.4, 128024), Ly6G (clone 1A8, 127618), CD11b (M1/70, 101212), NK1.1 (clone PK136, 108716), B220 (clone RA3-6B2, 103237), CD45 (clone 30F-11, 103138), CD8 (clone 341, 748879, BD Biosciences), CD29 (clone 30F-11, 103132), CD18 (clone M18/2, 101407), integrin β7 (clone FIB504, 321225), Ly6C (clone HK1.4, 128032), Ly6G (clone 1A8, 127648), CD8 (clone 53-6.7, 100712), CD4 (clone GK1.5, 100422), F4/80 (clone BM8, 123118), CD40 (clone 3/23 124610), PD-L1 (clone 10F.9G2, 124321) and CD206 (clone C068C2, 141731).

InVivoMAb anti-mouse CD29 (clone KMI6, BE0232), InVivoMAb anti-mouse/human integrin β7 (clone FIB504, BE0062) and InVivoMAb rat IgG2a isotype control, anti-trinitrophenol (clone 2A3, BE0089) antibodies were purchased from BioXCell.

For assessment of patient specimens, fluorophore-conjugated CD45 (clone HI30, 560777), CD33 (clone WM53, 562492), CD14 (clone M5E2, 558121), CD11c (clone 3.9, 748288), CD3 (clone SP34-2, 557757) and ITGB7 (clone FIB504, 555945) were purchased from BD Biosciences. CD19 (clone HIB19, 302218), CD56 (clone HCD56, 318332), Lox1 (clone 15C4, 358606), CD11b (clone ICRF44, 301332), CD66b (clone G10F5, 305114), HLA-DR (clone L243, 307638), CD29 (clone TS2/16, 303016), CD18 (clone CBR LFA-1/2, 366310), ITGB7 (clone FIB504, 121006), and CD68 (clone Y1/82A, 333814) were acquired from Biolegend.

### Mice

All animal experiments were approved by the Cleveland Clinic Institutional Animal Care and Use Committee (IACUC) and performed in accordance with the guidelines. Four-week-old C57BL/6 male and female mice (JAX Stock #000664) were purchased from the Jackson Laboratory as required and housed in the Cleveland Clinic Biological Research Unit Facility. Mice were intracranially injected at 4–8 weeks old with 25,000-30,000 GL261 or 15,000-30,000 SB28 cells in 5 μl RPMI null media into the left cerebral hemisphere 2 mm caudal to the coronal suture, 3 mm lateral to the sagittal suture at a 90° angle with the murine skull to a depth of 2.5 mm, using a stereotaxis apparatus (Kopf). Mice were monitored daily for neurological symptoms, lethargy and hunched posture that would qualify as signs of tumor burden.

### Adoptive transfer

For adoptive transfer, recipient mice were implanted with tumors as described above. A separate cohort of mice were implanted with tumors to obtain donor MDSCs for transfer. Femur and tibia from donor mice were flushed with 10 ml PBS and strained through 40 μm filter (Fisher Scientific). Cells were centrifuged at 400 *g* for 5 min and incubated with 1:25 diluted mouse FcR blocking reagent (Miltenyi Biotec, 130-092-575) in FACS Buffer (PBS, 5 mM EDTA and 2% FBS) on ice for 10 min. Samples were stained with a combination of 1:100 diluted CD11b, Gr-1 and Ly6G in the presence of FcR blocking reagent for 15 min on ice. Cells were resuspended at a concentration of 1-2 million/ml. mMDSCs (CD11b^+^Gr-1^+^Ly6G^-^) and gMDSCs (CD11b^+^Gr-1^+^Ly6G^+^) were sorted into FACS buffer using a BD FACSMelody (BD Bioscience). After centrifugation at 400 *g* for 5 min, the cells were resuspended in PBS at a concentration of 400,000 per 50 μl. In some experiments, sorted MDSCs were treated ex vivo with 100 μg/ml isotype control, anti-integrin β1 or anti-integrin β7 antibodies (BioXcell) for 1 hour on ice prior to transfer. Mice were anesthetized with isoflurane, and cells were retro-orbitally transferred using 27G needles (Exel).

### Immune profiling

GL261-bearing mice were adoptively transferred with MDSC subsets as described above, and mice were euthanized 3 days later. Cardiac blood was collected into EDTA-coated Safe-T-Fill^®^ micro capillary blood collection tubes (RAM Scientific). Samples were centrifuged at 1000 *g* for 10 min at 4°C to separate the serum, and cells were used for subsequent staining. Bone marrow was flushed from one femur and tibia in 5 ml PBS and passed through a 40 μm strainer to obtain single cells. Tumors were resected from the left hemisphere. From sham-injected controls, an equal volume of healthy brain tissue was removed. Tissue were mashed on a 40 μm strainer and washed with PBS before transferring into 96-well round-bottom plates (ThermoFisher Scientific). Samples were stained with 1:1000 diluted LIVE/DEAD Fixable Stains (ThermoFisher Scientific, L34962) in PBS for 10 minutes on ice. Following a wash step, cells were resuspended in FcR Blocking Reagent (Miltenyi Biotec) at a 1:25 dilution in PBS/2% BSA (Sigma-Aldrich) for 10 minutes on ice. Fluorophore-conjugated antibodies diluted 1:50 were added to suspensions, and cells were further incubated for 20 minutes on ice. Samples were washed with PBS/BSA and fixed overnight in eBioscience™ Foxp3/Transcription Factor Fixation Buffer. Samples were acquired with a Cytek Aurora (Cytek Biosciences) and analyzed using FlowJo (v10.7.2, BD). Statistically significant immune differences were determined by two-way ANOVA with Tukey’s correction for multiple comparisons. Individual immune populations were graphed separately.

### ATAC-Seq

mMDSCs and gMDSCs (2-4 million) were sorted from the bone marrow of sham-injected or GL261-implanted male and female mice. The experiment was performed in two biological replicates for each group, and cells from two mice were combined for each replicate. MDSCs were washed with cold PBS and counted on a Countess (Invitrogen). Pellets were resuspended in 1 ml of nuclei permeabilization buffer (PBS with 1 mM dithiothreitol, 5% bovine serum albumin, 0.2% IGEPAL-CA630 (Sigma Aldrich) and 1x cOmplete EDTA-free protease inhibitor (Roche)) and rotated for 10 min at 4°C as per the Ren et al. protocol (https://www.encodeproject.org/documents/4a2fc974-f021-4f85-ba7a-bd401fe682d1/). The nuclear suspension was filtered on a 30 μm CellTrics (Sysmex) and spun at 500 x g for 5 min at 4°C. The resulting pellet was resuspended in 50 μl tagmentation buffer (33 mM Tris-acetate pH 7.8, 66 mM potassium acetate, 11 mM magnesium acetate, and 16% N′N-dimethylformamide, in water). An aliquot of nuclear suspension was counted on a Countess. For each tagmentation reaction, 160,000 nuclei were mixed with 1 μl of Tagment DNA enzyme in a total of 20 μl and incubated for 30 min with 500 rpm at 37°C. Then 200 μl Buffer PB (Qiagen) and 10 μl sodium acetate (3M, pH 5.2) were added, and the sample was purified using a MinElute PCR purification kit (Qiagen) and eluted in 10 μl EB. Fragments were amplified by PCR following the Ren et al. protocol but using primers from Buenrostro et al. Supplementary Table 1, then purified using the MinElute PCR purification kit with elution in 40 μl EB (Buenrostro et al., 2013). Size selection was performed with PCRclean Dx beads (ALINE) by adding 160 μl EB to the 40 μl sample and adding 110 μl beads. After mixing and incubation for 5 min at room temperature, the tube was put on a magnetic stand and the supernatant transferred to a new tube, to which 190 μl beads was added. This suspension was mixed, incubated, and put on the magnetic stand, and this time the supernatant was discarded. Beads were washed twice with 70% ethanol, then the DNA was eluted from the beads with 20 μl EB. The indexed libraries were sequenced on a NextSeq high-output flowcell, paired-end, 75 cycles.

### ATAC-Seq data analysis

Cutadapt v1.9.1 (Martin, 2011) was used to remove paired-end adapter sequences and discard reads with a length less than 20bp. All FASTQs were aligned to the mm10 genome assembly (retrieved from http://hgdownload.cse.ucsc.edu/goldenPath/mm10/bigZips) using BWA-MEM v0.7.17-r1188 (Li, 2013) with default parameters in paired-end mode. Sequence alignment/map (SAM) output files were converted to binary (BAM) format, sorted, indexed, and PCR duplicates were removed using SAMtools v1.10 (Li et al., 2009). Peaks were detected with MACS v2.1.2 (Zhang et al., 2008) with --format=BAMPE. DeepTools v3.2.0 (Ramirez et al., 2016) was used to generate RPGC-normalized bigWig tracks with 50 bp bin sizes of the final sample BAM files and aggregate bigWig files were generated by averaging the ATAC-Seq signals across either all gMSDC or mMSDC samples. Libraries were assessed for quality using ChIPQC (Carroll et al., 2014) and visualized on the Integrative Genomics Viewer (Thorvaldsdottir et al., 2013). Peak lists were filtered to remove all peaks overlapping ENCODE blacklisted regions (mm10 blacklist file v2 downloaded from https://sites.google.com/site/anshulkundaje/projects/blacklists).

### Identification of differential open chromatin regions

ATAC-Seq peaks called across all samples were filtered for significance (peaks with Benjamini-Hochberg corrected p-values (q-val) > 0.001 were excluded), combined together, overlapping peaks merged, and read depth for each peak region across samples determined using BEDTools v2.17.0 (Quinlan and Hall, 2010), generating a count matrix. A total of 89,775 merged peaks were used for differential testing. Peaks were tested for differential expression by cell type using DESeq2 v1.32.0 (Love et al., 2014) after controlling for sex and tumor status and stratifying for peaks with mean normalized counts > 50. Differential open chromatin regions were designated as gained or lost by a positive or negative 2-fold change in ATAC signal between mMDSC and gMDSC samples at q-val < 0.001.

### Gene mapping and ontology analysis

The functional enrichment analysis software Genomic Regions Enrichment of Annotations Tool (GREAT) (McLean et al., 2010) was used to map genes to peaks and to identify ontology terms associated with differential peaks. The following statistical test thresholds were used to identify significant ontology terms: binomial fold-enrichment > 2.0, binomial FDR Q-value < 0.05, and hypergeometric FDR Q-value < 0.05.

### Single-cell expression analysis

Intra-study normalized matrices of non-tumor cells for each of the four studies were downloaded from GigaScience GigaDB database (http://gigadb.org/dataset/100794) (Caruso et al., 2020). Data was combined from 4 studies containing 50 GBM samples and 9097 non-tumor cells in total. the distribution of samples is as follows: Darmanis et al., GEO Accession GSE84465, 2498 nontumor cells, n=4 patients, Smart-seq2 technology; Yuan et al., GEO Accession GSE103224, 4194 non-tumor cells, n=10 patients, Proprietary microwell technology; Neftel et al., GEO Accession GSE131928, 1067 non-tumor cells, n=28 patients, Smart-seq2 technology; Yu et al., GEO Accession GSE117891, 1338 non-tumor cells, n=8 patients, STRT-seq technology. Basic analysis and visualization of the scRNA-seq data were performed with the Seurat R package (v.4.0.2) in R (v.3.3.4). Matrices were processed by SCTransform() and IntegrateData() functions to achieve inter-study normalization and integration (https://satijalab.org/seurat/articles/integration_introduction.html#performing-integration-on-datasets-normalized-with-sctransform-1). The default value was used when running the functions (Hafemeister and Satija, 2019; Hao et al., 2021). The myeloid cell population was defined based on high ITGAM expression, and subsets were further discriminated based on relative expression profiles of CD84, CD33, ITGAM (CD11b), CD14, OLR1 (LOX-1), CEACAM8 (CD66) and HLA-DR (**Fig. S2B**).

### Analysis of patient tumors

Five GBM specimens were collected by the Rose Ella Burkhardt Brain Tumor and Neuro-Oncology Center in accordance with the Cleveland Clinic Institutional Review Board (IRB 2559). Tumors were minced and processed following the instructions for the human tumor dissociation kit (Miltenyi Biotec, 130-095-929). After the addition of enzyme mix in 5 ml null RPMI, samples were digested in a Miltenyi dissociator using the 37_h_TDK_1 program. Cells were treated with RBC Lysis Buffer (Biolegend) at room temperature for 5 minutes. Samples were stained with LIVE/DEAD Fixable Stains for 10 minutes on ice and incubated with FcR Blocking Reagent for 15 minutes on ice. Staining with fluorophore-conjugated CD45, CD33, CD14, CD11c, CD3, CD19, CD56, Lox1, CD11b, CD66, HLA-DR, CD19, CD28 and integrin β7 antibodies was performed in Brilliant Stain Buffer (BD Biosciences) for 20 minutes on ice. Cells were fixed overnight in eBioscience™ Foxp3/Transcription Factor Fixation Buffer. CD68 staining was performed in eBioscience™ Foxp3/Transcription Factor Permeabilization Buffer with 20 minutes of incubation at room temperature. Samples were acquired with a BD LSR Fortessa (BD Biosciences).

### Macrophage polarization

Sorted mMDSCs (50,000) were stimulated with 50 ng/ml recombinant mouse M-CSF (Biolegend) in Iscove’s Modified Dulbecco’s Medium (IMDM, Media Preparation Core) with 1% Pen/Strep and 20% FBS in the presence of 100 ng/ml anti-integrin β1 blocking antibody for 7 days. Macrophages were removed by Accutase treatment, stained with LIVE/DEAD Fixable Stain for 10 minutes on ice and incubated with FcR Blocking Reagent for 15 minutes on ice. Samples were stained with a cocktail of 1:100 diluted CD68, F4/80, I-A/I-E, CD40, CD206 and PD-L1 antibodies for 20 minutes on ice before analyzing with a BD LSR Fortessa.

### Statistical analysis

GraphPad PRISM (Version 9, GraphPad Software Inc.) software was used for data presentation and statistical analysis. Two-way ANOVA and paired t-tests were used for comparison of differences among sample groups. The Gehan-Breslow-Wilcoxon test was used to analyze survival data. The specific statistical method employed for individual data sets is listed in the figure legends.

## Supplementary Figure Legends

**Supplementary Figure 1:**
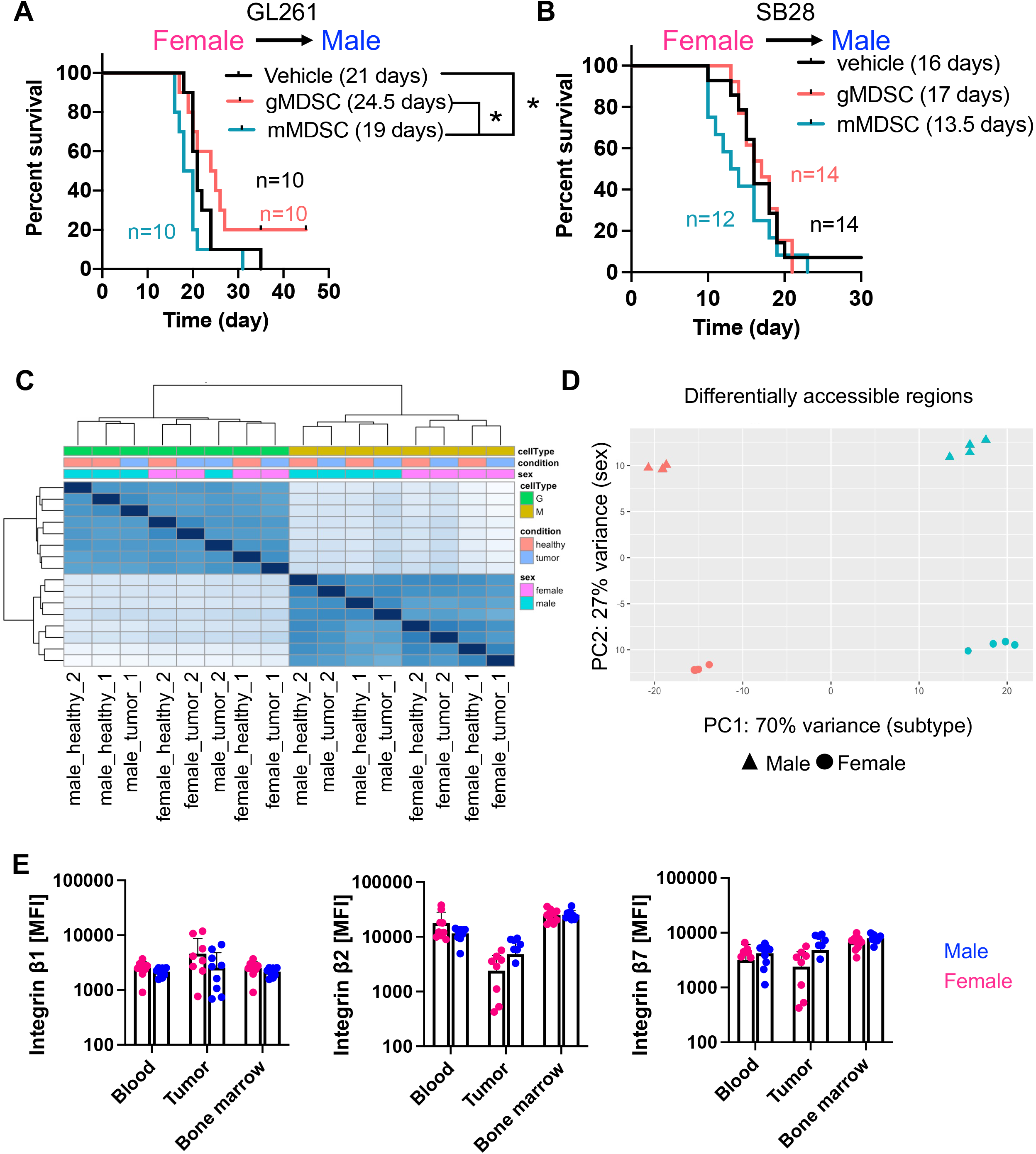
Sex has a minimal impact on mMDSC behavior, epigenetic profile and integrin expression. Male C57BL/6 mice were implanted with 25,000 GL261 or 10,000 SB28 cells. Seven (SB28) or 14 (GL261) days post-tumor implantation, mice were adoptively transferred with 400,000 mMDSCs or gMDSCs isolated from the bone marrow of female mice with matching tumors by retro-orbital injection. Kaplan-Meier curves depicting survival of **(A)** GL261- or **(B**) SB28-bearing mice post adoptive transfer. n=10-14 mice/group from 2-3 independent experiments. * p<0.05 as assessed by Wilcoxon test. **(C)** Clustering analysis demonstrating the impact of cell type, tumor type and sex on the chromosome accessibility profile. **(D)** Principal component analysis (PCA) depicting the relative contribution of cell type (major) and sex (minor) on differential chromosome accessibility. **(E)** Mean fluorescence intensity of surface ITGB1, ITGB2 and ITGB7 expression on male versus female mMDSCs from n=9-10 tumor (GL261/SB28)-bearing mice.

**Supplementary Figure 2:**
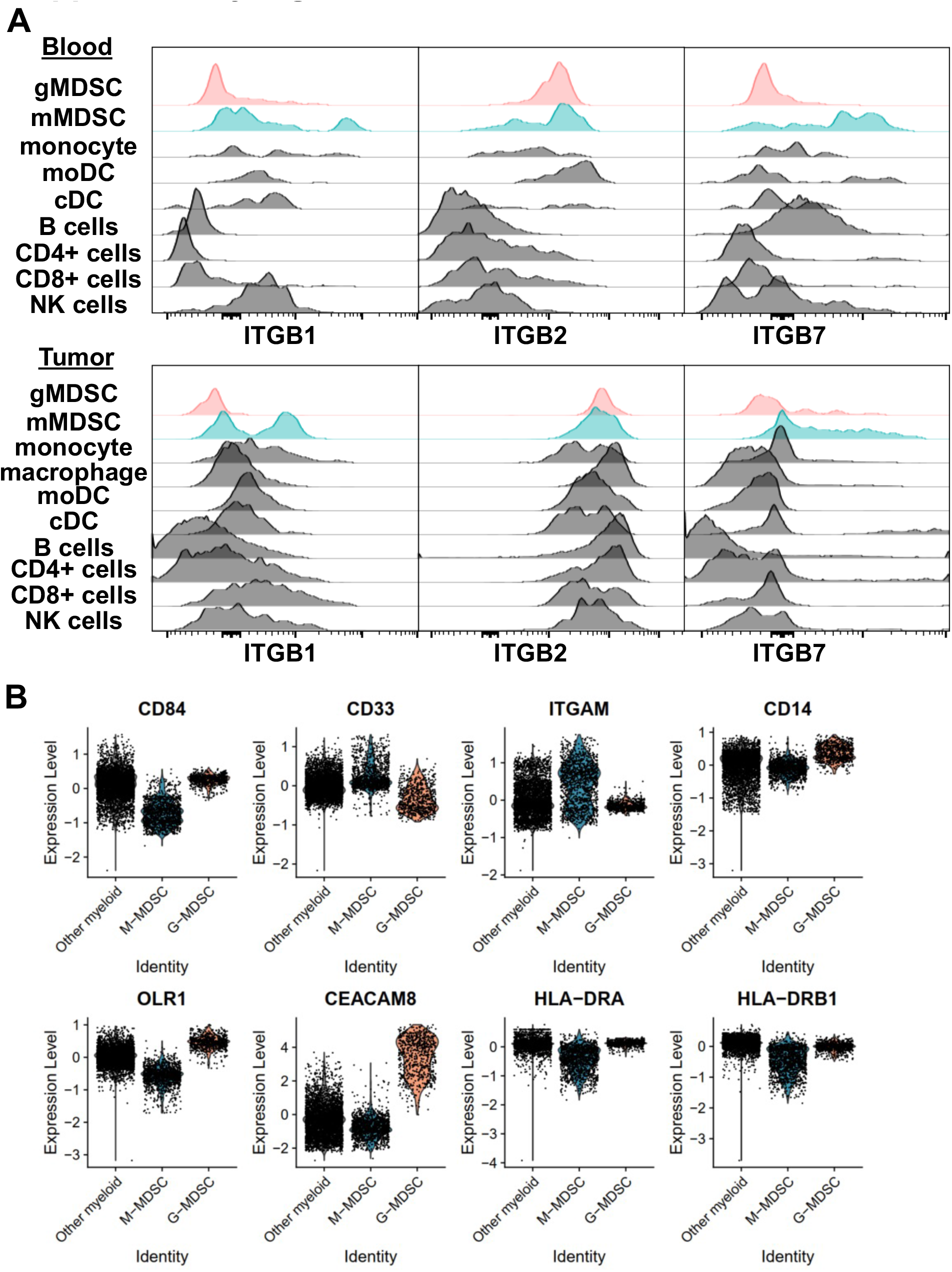
Integrin β1 and β7 are expressed at high levels by monocytic cells. C57BL/6 mice were implanted with 25,000 GL261 cells, and surface integrin levels were determined from circulating and tumor-infiltrating immune cell populations. **(A)** Histograms depicting relative expression of ITGB1, ITGB2 and ITGB7 across gMDSCs, mMDSCs, monocytes, macrophages, myeloid DCs, conventional DCs, B cells, CD4+ T cells, CD8+ T cells and NK cells in blood (top) or tumors (bottom). (**B)** Expression levels of CD84, CD33, ITGAM (CD11b), CD14, OLR1 (LOX-1), CEACAM8 (CD66) and HLA-DR were used to identify mMDSCs and gMDSCs in the tumors.

**Supplementary Figure 3:**
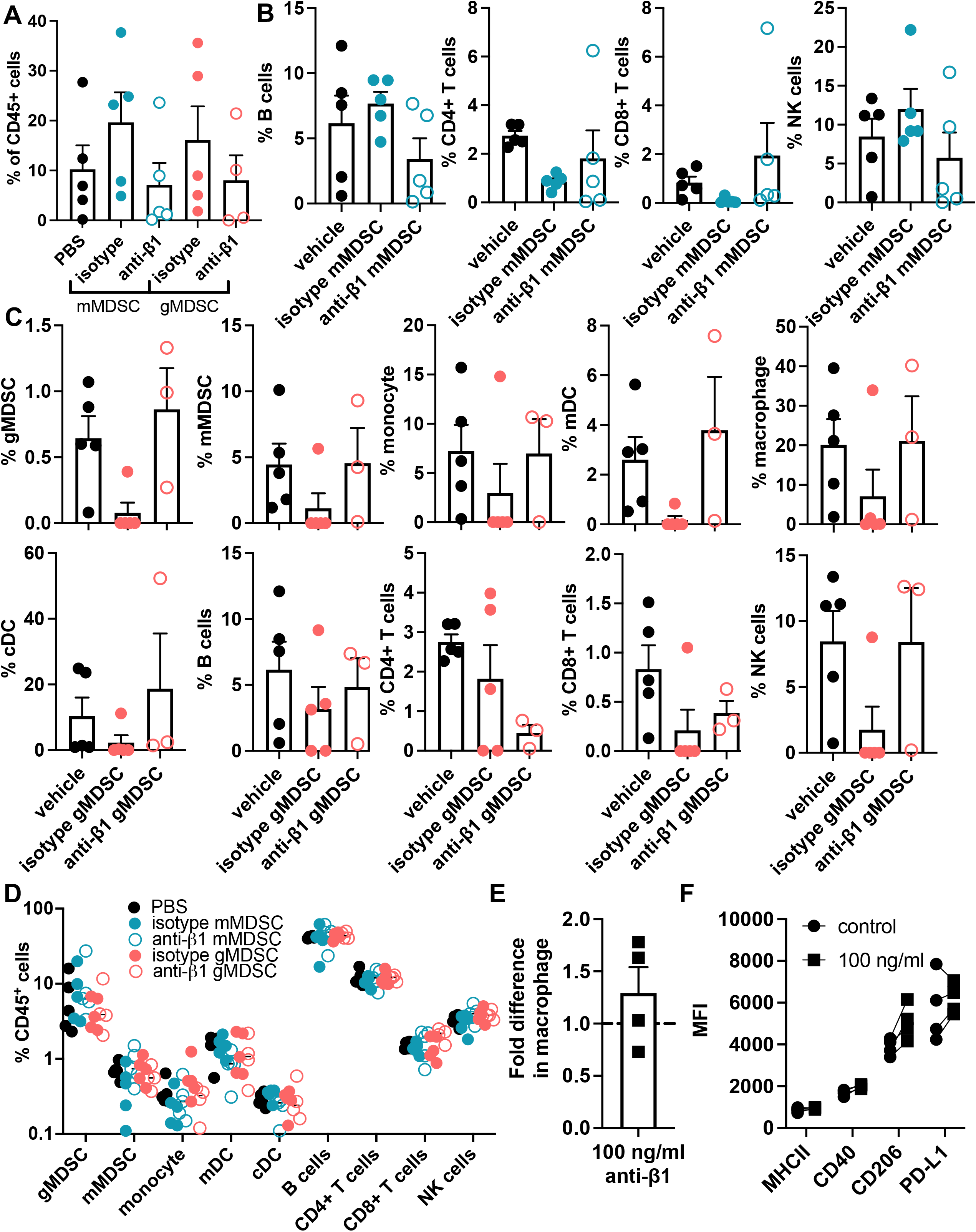
Myeloid blockade of integrin β1 does not impact tumor lymphocyte infiltration or systemic immunity. C57BL/6 male mice were implanted with 25,000 GL261 cells and adoptively transferred with 400,000 MDSC subsets treated with isotype control antibody or anti-integrin β1 blocking antibody for 1 hour. Animals were euthanized 3 days posttransfer, and immune populations were analyzed from blood and tumors. (A) Frequency of leukocytes in tumors of mice adoptively transferred with mMDSCs or gMDSCs. (B) Percentage of B cells, CD4+ T cells, CD8+ T cells and NK cells within the tumor-infiltrating leukocyte population of mice adoptively transferred with isotype- or anti-integrin β1-treated mMDSCs. (C) Percentage of gMDSCs, mMDSCs, monocytes, macrophages, myeloid DCs, conventional DCs, B cells, CD4+ T cells, CD8+ T cells and NK cells within the tumor-infiltrating leukocyte population of mice adoptively transferred with isotype- or anti-integrin β1-treated gMDSCs. (D) Frequency of immune populations in the peripheral circulation following adoptive transfer of mMDSCs and gMDSCs. n=3-5/group. mMDSCs were stimulated in vitro with M-CSF for 7 days in the presence of 100 ng/ml anti-integrin β1 blocking antibody. The number and phenotype of macrophages were assessed with flow cytometry. (E) Relative number of macrophages in anti-integrin β1 cultures compared to isotype control. (F) Expression levels of MHCII, CD40, CD206 and PD-L1 in the resultant macrophages. n=2 male and 2 female.

